# NanoBlocks: creating fluorescent biosensors from affinity binders using competitive binding

**DOI:** 10.1101/2025.10.28.685015

**Authors:** Silke Denis, Vincent Van Deuren, Desiree I. Frecot, Ulrich Rothbauer, Shehab Ismail, Alison G. Tebo, Peter Dedecker

## Abstract

Genetically-encoded fluorescent biosensors have revolutionized our understanding of complex systems by permitting the *in situ* observation of chemical activities. However, only a comparatively small set of chemical activities can be monitored, largely due to the need to identify protein domains that undergo conformational and/or association changes in response to a stimulus. Here, we present a strategy that can convert ‘simple’ affinity binders such as nanobodies into biosensors for their innate targets by introducing a peptide sequence that competes for the binding site. We demonstrate proof-of-concept implementations of this ‘NanoBlock’ design, developing sensors based on the ALFA nanobody and on the PDZ domain of Erbin. We show that these sensors can reliably detect their targets *in vitro*, in mammalian cells, and as part of fluorescence-activated cell sorting (FACS) experiments. In doing so, our strategy offers a way to strongly expand the range of cellular processes that can be probed using fluorescent biosensors.

## Introduction

Fluorescence microscopy is an invaluable tool to unravel the myriad of processes occurring in biological systems. Fluorescent biosensors have vastly expanded the scope of what can be observed in living systems because they can visualize the spatiotemporal dynamics of enzymatic activities (*1*), ions (*2, 3*), neurotransmitters (*4, 5*) and other cellular biochemistry. A variety of biosensor architectures have been developed for different purposes and targets (*6*). However, despite their transformative potential, biosensors face a critical bottleneck: the limited availability of protein domains that can serve as molecular switches to detect new targets of interest.

Current biosensor designs all rely on stimulus-sensitive proteins that change conformation upon target binding, thereby altering the optical properties of a fluorescent reporter domain (*7*). In domain insertion designs, the fluorescent domain is inserted into the sensing domain or vice versa to tightly couple target recognition and fluorescent signal. Alternatively, two sensing moieties that change association when the sensed stimulus is present can be fused to the termini of a fluorescent domain, as is the case for the extremely successful genetically-encoded calcium indicators (GECIs) (*8*–*10*), kinase/phosphatase reporters (*1, 11*–*13*), and others. However, both strategies share a fundamental constraint in their requirement for protein domains that can act as molecular switches (*7, 14*), thus limiting the range of stimuli that can be visualized since few such domains are known. While sensors based on molecular translocation can circumvent this requirement, these require the availability of domains that enable this translocation behavior, are often associated with slow response times, cannot easily report on highly localized changes in stimulus activity, and require high-resolution imaging, which may be difficult to achieve in optically large samples (*14*).

While developing novel, conformationally dynamic sensing domains remains difficult, it is comparatively much easier to discover or develop affinity binders that ‘simply’ bind their targets but do not undergo significant binding-induced conformational changes. For example, numerous single-domain affinity binder scaffolds have been created, including DARPins (*15*), affibodies (*16*), and nanobodies (*17*). We reasoned that the existence of a general methodology to transform such artificial or natural binders into biosensors could broadly expand the range of stimuli that can be sensed. Several efforts have aimed to deliver such a capability, though so far the resulting systems still have some drawbacks like slow response times (*18*), limited choice of affinity binders (*19*), incompatibility with expression in live mammalian cells (*20, 21*) or large, bimolecular constructs that need extensive computational engineering (*22, 23*).

In this contribution, we present a novel sensor architecture that offers a more generic way to create unimolecular fluorescent biosensors from affinity binders. Our strategy relies on the introduction of a peptide that competes for the binding site of the binder, thus realizing a large conformational change when the stimulus is present. Our design results in quick-responding, live-cell compatible sensors that we call ‘NanoBlock’ sensors. Overall, our architecture can strongly expand the range of processes that can be visualized *in situ*.

## Results and disscussion

### NanoBlock biosensor architecture

The overall concept underlying our sensors is shown in Figure 1a. Briefly, we make use of an affinity binder such as a nanobody fused to a circularly permuted fluorescent protein (cpFP). At the other terminus of the cpFP, we introduce a peptide that weakly binds to the affinity binder, such that it is bound when the ligand of the binder is absent. Addition of the high-affinity ligand will outcompete this peptide, resulting in its displacement and a large conformational change transmitted to the cpFP. The transduction of this conformational change to the cpFP would then create a functional biosensor with fluorescence properties dependent on the presence of the target. Because one could think of the peptide as a ‘binding’ or perhaps a ‘blocking’ motif, we named molecules created in this way ‘NanoBlocks’.

**Figure 1:**
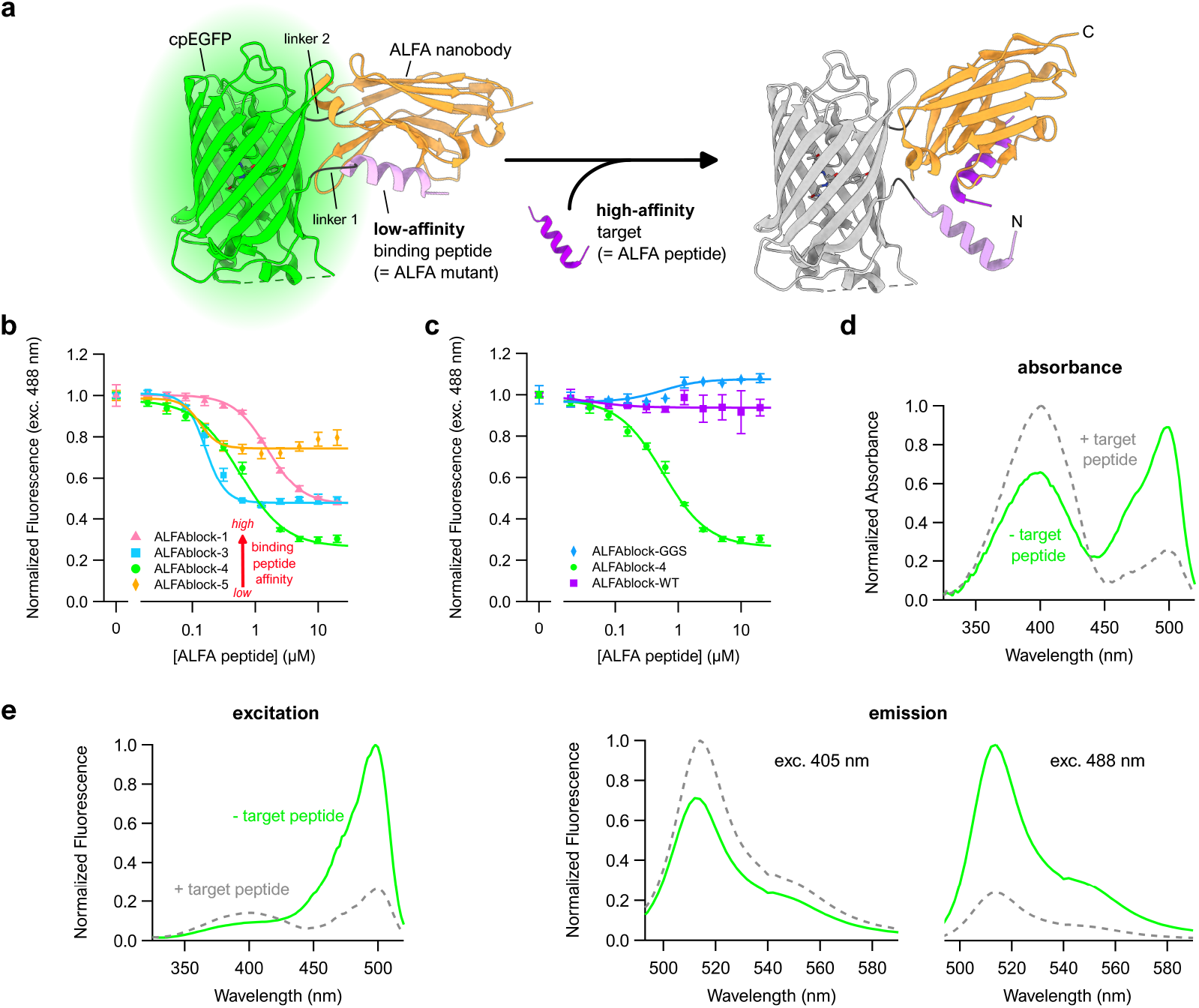
ALFAblocks – NanoBlock biosensors that recognize the ALFA-tag. **a)** Schematic representation and proposed response mechanism of the ALFAblocks. To depict the sensor, parts of crystal structures 6I2G and 3WLC (PDB) were assembled manually using UCSF ChimeraX (*29*). **b, c)** ALFA-tag peptide titration curves of ALFAblocks containing binding peptides with different affinities **(b)** or control binding peptides **(c)**. Solid lines represent Hill equation fits, data points represent mean ± s.d., n=3. Purified sensor at 100 nM or 500 nM (GGS). **d)** Absorbance and **e)** fluorescence spectra of ALFAblock-4, with and without a saturating concentration of ALFA-tag peptide. Mean of n=2.

The main advantage of this design is that it is generic in nature since replacement of the affinity binder-binding peptide pair will generate a biosensor that responds to a different stimulus. The main disadvantage is that a suitable peptide that competes for the stimulus binding site must be available. Fortunately, the binding affinity of this peptide should be sufficiently low that it is outcompeted by the natural target, which reduces the design requirements.

The concept of competitive binding to create biosensors has been previously used in the Snifit sensors (*24, 25*), in the Ras activity sensor RasAR (*26*) and in the LOCKR switch platform (*22, 23*). However, in comparison to these already existing strategies, our NanoBlock design stands out because it is completely genetically encoded, its architecture is relatively compact, the readout is conceptually simple, and the design offers enhanced generalization to other readouts or reporters.

### Development and characterization of ALFAblocks

We sought to develop a proof-of-concept implementation of our idea via the creation of ‘ALFAblocks’. These are NanoBlocks based on the ALFA nanobody (NbALFA), which binds a peptide known as the ALFA-tag (*27*). The ALFA-tag is a 13 amino acid helical peptide that binds with sub-nanomolar affinity (*K*_*D*_ = 165 ± 2 pM, Table 1) and that is reported to not interfere with the folding or cellular localization of a target to which it is fused. To obtain a prototype ALFAblock construct, we combined circularly-permuted EGFP (cpEGFP) with an N-terminal binding peptide and a C-terminal NbALFA (Figure 1a). The cpEGFP variant and the linkers that connect the different domains were taken from the calcium sensor GCaMP6s (*28*).

**Table 1:** ALFA-like binding peptide properties and sensor parameters of the corresponding purified ALFAblocks.

| ALFA-like peptide | WT | 1 | 3 | 4 | 5 | GGs |
| --- | --- | --- | --- | --- | --- | --- |
| $K_{\text{pep}}$ (nM)<sup>a</sup> | 0.165 ± 0.002 | 3.05 ± 0.0141 | 3.93 ± 0.0265 | 68.7 ± 1.66 | 111 ± 5.29 | ND |
| $k_{\text{on}}$ ($10^5 \text{M}^{-1} \text{s}^{-1}$)<sup>a</sup> | 0.68 ± 0.0027 | 1.51 ± 0.0067 | 2.30 ± 0.0152 | 5.80 ± 0.134 | 5.71 ± 0.259 | ND |
| $k_{\text{off}}$ ($10^{-3} \text{s}^{-1}$)<sup>a</sup> | 0.013 ± 0.00015 | 0.462 ± 0.00075 | 0.906 ± 0.00116 | 39.8 ± 0.286 | 63.2 ± 0.967 | ND |
| ALFABlock- | WT | 1 | 3 | 4 | 5 | GGs |
| $\Delta F/F^b$ | 0.09 | 1.10 | 1.05 | 2.30 | 0.26 | 0.08 |
| $K'_D$ ($\mu\text{M}$)<sup>c</sup> | NA | 1.53 ± 0.07 | 0.11 ± 0.01 | 0.53 ± 0.03 | 0.08 ± 0.02 | NA |
| Hill Coefficient<sup>d</sup> | NA | 1.60 ± 0.10 | 2.76 ± 0.22 | 1.21 ± 0.08 | 2.96 ± 0.75 | NA |
| $C_{\text{eff}}$ binding peptide ($\mu\text{M}$)<sup>e</sup> | NA | 28.28 | 2.62 | 220.60 | 53.71 | NA |
| Relative Brightness<sup>f</sup> | 0.22 | 1.12 | 0.47 | 1.00 | 0.33 | 0.39 |
| Time Constant (min)<sup>g</sup> | NA | 16.19 | 0.75 | 4.19 | 0.37 | NA |

Four potential binding peptides were tested (1, 3, 4, and 5), all ALFA-like peptides with varying but lower affinities for NbALFA (Table 1 and Supplementary Figure S1). All of these constructs (ALFAblock-1, -3, -4 and -5) showed a clear decrease in fluorescence intensity upon addition of the ALFA-tag peptide (Figure 1b), indicating that they were indeed functional biosensors for the presence of the ALFA-tag. Our proposed mechanisms suggest that this reduction in fluorescence intensity arises from a loss of the interaction between the nanobody and binding peptide, consistent with what is seen for structurally similar sensors such as GCaMP sensors, where Ca^2+^ binding induces domain association and an increase in fluorescence (*28*).

We created two control sensors to further validate our proposed sensing mechanism. In the first control (ALFAblock-GGS), we replaced the binding peptide with a GGSGGSG sequence, thus eliminating the affinity between the binding peptide and nanobody. In the second control (ALFAblock-WT), we replaced the ALFA-like binding peptide with the wild type ALFA-tag sequence. Accordingly, we expected that the GGS-control would always be in the ‘open’ state, while the WT-control would always be in the ‘closed’ state due to the very high affinity of the wild-type ALFA-tag. Indeed, neither of the control ALFAblocks responded significantly to the addition of free ALFA-tag peptide (Figure 1c), consistent with a sensing mechanism that involves displacement of the binding peptide from the affinity binder.

We next quantified the observed performance by calculating the fluorescence contrast (Δ*F*/*F*), the fluorescence brightness, and the apparent dissociation constant *K*_D_, (Table 1). We also constructed a theoretical model that allowed us to express the overall NanoBlock binding affinity to its target as a function of the nanobody affinities for its natural target and for the binding peptide (Supplementary Note S1). Briefly, we find that the apparent dissociation constant of a NanoBlock is equal to 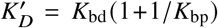, where *K*_bd_ is the dissociation constant for the natural target and *K*_bp_ is the dissociation constant for the binding peptide when both are present in a single construct.

*K*_bp_ is determined by the natural affinity of the binding peptide to the nanobody (given by its dissociation constant *K*_pep_ when both are free in solution) as well as the local enrichment and possible steric effects that arise via their fusion in a single construct. Our theoretical model allows us to express this via *K*_bp_ = *K*_pep_ /*C*_eff_, where *C*_eff_ is the effective concentration of the binding peptide. Briefly, we expect *C*_eff_ to be equal to the total concentration of the sensor if the binding peptide is equally likely to bind via an intramolecular as via an intermolecular (dimer) configuration. Higher values for *C*_eff_ show increased intramolecular binding and accordingly reduced steric hindrance to such binding. This framework provides a deeper explanation for the different values observed in our experiments. In particular, we find that the effective concentrations are much higher than the sensor concentration (0.1 µM) for all of the constructs used, showing that these NanoBlocks indeed function via intramolecular association of the binder and peptide moieties. We do find particularly low values for the construct making use of peptide 3, which may arise from sequence-specific intramolecular interactions that lower the availability of the peptide for the nanobody.

Time-resolved measurements revealed that the sensor responses to ALFA peptide addition occurred over a few minutes or less, with faster time constants for lower-affinity binding peptides (Supplementary Figure S2 and Table 1). These response times could likely be improved by further reducing the affinity of the binding peptide, but are still considerably faster than other nanobody-based biosensors like the ATOM sensors (*18*), which need several hours to respond due to their dependence on chromophore maturation.

We selected ALFAblock-4 for further characterization since it stood out in terms of brightness and dynamic range. Spectroscopic analysis of this construct revealed two absorption and excitation peaks at approximately 400 nm and 500 nm, which increased and decreased after ALFA-tag addition, respectively (Figure 1d-e). As is commonly observed, these peaks correspond to the neutral (400 nm) and anionic (500 nm) chromophores with both giving rise to a single emission peak around 515 nm through excited state proton transfer (ESPT) from the excited neutral form of the chromophore (*30*). Similar to other sensors based on cpFPs, the redistribution of the chromophore population from anionic to neutral is the main force behind the observed fluorescence contrast upon target binding (*31*). As a consequence, NanoBlock sensing can be performed using excitation at either 488 or 405 nm, leading to either a negative contrast (fluorescence decrease) and Δ*F* /*F* = 2.30 for excitation using 488 nm, and a positive contrast with Δ*F /F* = 0.47 for excitation using 405 nm (Supplementary Figure S3). This also permits an excitation-ratiometric readout when the 488 nm-excited fluorescence is divided by the 405 nm-excited fluorescence. This results in a readout that is in principle independent of the sensor concentration, with a ratiometric dynamic range Δ*R*/*R* = 3.47 (negative contrast). Interestingly, ALFAblocks-1, -3 and -5 do not respond to the ALFA peptide when excited with 405 nm-light only (Supplementary Figure S2 and S3).

pH titrations showed that ALFAblock-4 in the absence of free ALFA peptide has a pKa of 7.42 ± 0.02 (exc. 488 nm), which increases to 7.98 ± 0.02 when a saturating amount of peptide is added (Supplementary Figure S4). The effect of pH is observable for both 405 and 488 nm excitation and also exerts a marked effect on the ratiometric response. This pKa near physiological pH complicates the robustness of the sensor readout since a change in cellular pH could generate a response that is mistaken for a change in the activity of the sensed target, though our observation is in line with the pH dependence observed for many other GCaMP-like sensors (*3, 5, 10, 32*).

### ALFAblock optimization and mammalian cell expression

We next sought to improve the sensing properties of ALFAblock-4. We chose to optimize the linkers between the different sensor moieties, a well-known and successful approach for the development of single FP-based biosensors (*1, 7, 33*). We conducted a bacterial lysate screen of about 1500 mutants expressing varying linker lengths (1-3 aa (amino acids)) and compositions (Supplementary Figure S5a). We chose to randomize the linker residues closest to the cpEGFP since these were shown to have the most influence on sensor performance (*7*). Our choice of linker lengths is customary for single FP-based sensors and should ensure a short enough linker to efficiently transduce conformational changes while preserving correct folding of all domains. Both linkers were optimized simultaneously.

The screen yielded several variants with improved dynamic range, response kinetics, and/or 488 nm-excited fluorescence brightness (Supplementary Figure S5b-c). We included this last criterion because higher fluorescence intensity in this screen could indicate either improved intrinsic brightness or improved expression, folding and/or maturation, all desirable properties in a biosensor. We even identified ‘turn-on’ variants in which target addition resulted in an increase of fluorescence, although the dynamic ranges were considerably lower compared to their best turn-off counterparts.

We found that the N-terminal linker displayed the strongest influence on the sensor properties. For example, an N-terminal linker of one amino acid typically results in a turn-on sensor. A possible explanation could be that this very short linker between binding peptide and cpEGFP causes part of the cpEGFP β-barrel to unravel in the ALFA-tag-free, ‘closed’ conformation, increasing solvent accessibility and thus protonation of the chromophore. When the ALFA-tag binds and the binding peptide is displaced from the nanobody, the binding peptide and linker become more flexible and could allow full refolding of the barrel and deprotonation of the chromophore (*7, 34*). Another observation is that an N-terminal linker of 2 or 3 amino acids in combination with an aromatic residue closest to the cpEGFP gives rise to a turn-off sensor with high Δ*F /F* (between 3.1 and 7.4 in bacterial lysates). Crystal structures of the Ca^2+^-bound GCaMP2 (PDB 3EK4) and GCaMP6m (PDB 3WLD) sensors reveal that the corresponding residue points towards the chromophore, suggesting that it plays in role stabilizing the hydrogen bond network that keeps the chromophore in its deprotonated state (*34, 35*).

To evaluate the performance of the ALFAblocks in mammalian cells, the most promising linker mutants in terms of dynamic range and/or fluorescence intensity were expressed in the cytoplasm of HeLa cells (Supplementary Figure S5d). However, the sensor fluorescence was not distributed homogeneously in the majority of cells but instead aggregated in fluorescent puncta. Interestingly, double transfection with an ALFA-tagged mCherry protein without functional chromophore (‘dark’ mCherry) avoided the formation of puncta, possibly increasing solubility through the interaction between NbALFA and ALFA-tag. The propensity of cells to form such puncta was reduced but not eliminated by fusing the protein to solubility tags (maltose-binding protein, B1 domain of streptococcal protein G, a non-fluorescent variant of mCherry) and the mammalian expression of ALFAblocks was therefore not further pursued.

### Development and characterization of ePDZblocks

We next sought to explore the generic nature of our concept by applying it to a different class of affinity binder. We selected the PDZ domain of Erbin, a human signaling scaffold protein. Phage display experiments have identified peptides that bind with differing affinities to this domain (*36*), including competition ELISA IC_50_’s of 0.6, 14, and 480 µM. We refer to these peptides as ‘0.6’, ‘14’, and ‘480’, respectively.

We created two ‘ePDZblocks’ by incorporating the ePDZ domain and the 14 and 480 binding peptides into the GCaMP6s framework. Since ePDZ specifically interacts with the C-terminal carboxylate of these peptides, the binding peptide was placed C-terminally of cpEGFP while ePDZ was moved to the N-terminus of the NanoBlock, which is the inverse of the ALFAblocks (Figure 2a). Two control sensors were created as well, in which the binding peptides are expected to have no (ePDZblock-GGS) or a high affinity (ePDZblock-0.6) for the ePDZ domain.

**Figure 2:**
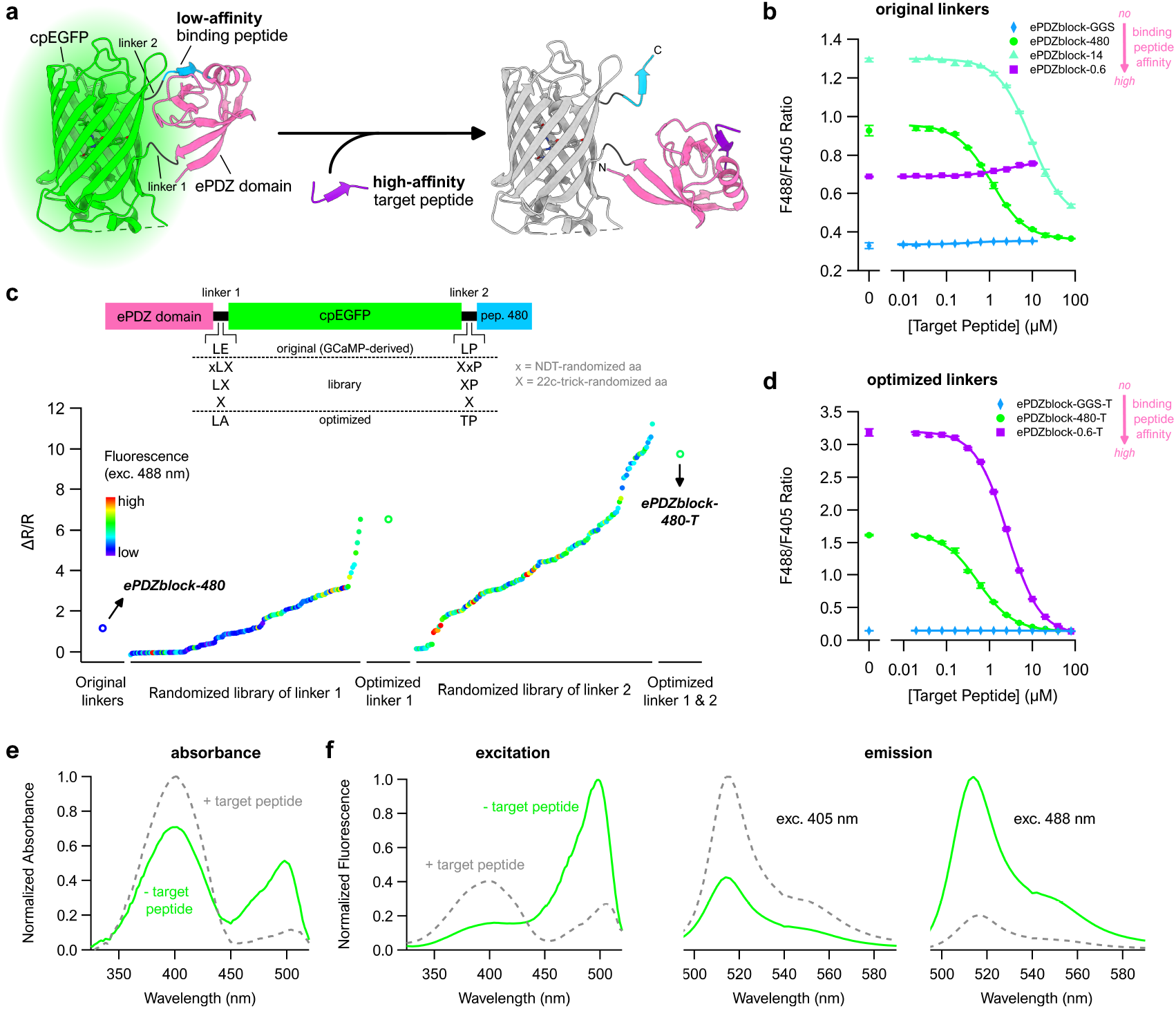
ePDZblocks – NanoBlock biosensors that recognize the 0.6 target peptide. **a)** Schematic representation and proposed response mechanism of the ePDZblocks. To depict the sensor, parts of crystal structures 6Q0N and 3WLC (PDB) were assembled manually using UCSF ChimeraX (*29*). **b)** Target peptide titration curves of ePDZblocks with different binding peptides. Sensors contain the original, GCaMP-derived linkers. Solid lines represent Hill equation fits, data points mean ± s.d., n=3. Purified sensor at 100 nM. **c)** Linker optimization of ePDZblock-480 through bacterial lysate screens. The first screen focused on linker 1 and covered 149 colonies, each containing a linker 1 variant. Once an optimized sequence was selected for linker 1, this was kept constant and in a second library, linker 2 was randomized. 166 more colonies were screened. Each dot represents one screened linker variant, ranked by increasing ratiometric dynamic range (negative contrast). **d)** Target peptide titration curves of ePDZblocks with different binding peptides. Sensors contain the optimized linkers from the bacterial lysate screen. Solid lines represent Hill equation fits, data points mean ± s.d., n=3. Purified sensor at 100 nM. **e)** Absorbance and **f)** fluorescence spectra of ePDZblock-480-T, with and without a saturating concentration of target peptide. Mean of n=3.

Titration of the four constructs with the high-affinity 0.6 peptide revealed that we had indeed created working biosensors against ePDZ binding partners (Figure 2b, Table 2 and Supplementary Figure S6a). Both ePDZblock-14 and -480 show decreased fluorescence upon target addition (Δ*F /F* = 1.64 for ePDZblock-14 and 1.33 for ePDZblock-480, exc. 488 nm), while ePDZblock-GGS shows no response to the target. ePDZblock-14 does not respond to the target when excited with 405 nm-light, while ePDZblock-480 is very suitable for use with an excitation-ratiometric approach and has an increased ratiometric dynamic range (Δ*R /R* = 1.54, negative contrast). The ePDZblock-0.6 fluorescence increases slightly under both 488 and 405 nm-excitation, but the ratiometric signal stays relatively unresponsive. The apparent sensor affinity of the ePDZblocks is inversely proportional to the affinity of the binding peptide for ePDZ (Table 2), consistent with our model (Supplementary Note S1) and the behavior observed on the ALFAblocks.

**Table 2:** ePDZ binding peptide properties and sensor parameters of the corresponding purified ePDZblocks.

| **Binding Peptide** | **0.6** | **14** | **480** | **GGs** |
| --- | --- | --- | --- | --- |
| Sequence | TGWETWV | QPVDSWV | ATQITWV | GGSGGSG |
| Competition ELISA IC<sub>50</sub> (μM)<sup>a</sup> | 0.6 | 14 | 480 | ND |
| K<sub>pep</sub> (μM) | 0.24 (37) | 2.2 (38) | ND | ND |
| **ePDZblock** | **0.6** | **14** | **480** | **GGs** |
| ΔR/R<sup>b</sup> | 0.10 | 1.42 | 1.54 | 0.08 |
| K'<sub>D</sub> (μM)<sup>c</sup> | NA | 9.20 ± 0.26 | 1.07 ± 0.02 | NA |
| Hill Coefficient<sup>d</sup> | NA | 1.13 ± 0.02 | 1.10 ± 0.01 | NA |
| C<sub>eff</sub> binding peptide (μM)<sup>e</sup> | NA | 82.13 | 1660 | NA |
| Relative Brightness<sup>f</sup> | 0.70 | 1.21 | 1.00 | 0.67 |
| Time constant (s)<sup>g</sup> | NA | 24 | 20 | NA |
| **ePDZblock-T** | **0.6** | **14** | **480** | **GGs** |
| ΔR/R<sup>b</sup> | 22.65 | ND | 10.38 | 0.02 |
| K'<sub>D</sub> (μM)<sup>c</sup> | 2.59 ± 0.03 | ND | 0.49 ± 0.01 | NA |
| Hill Coefficient<sup>d</sup> | 1.16 ± 0.01 | ND | 1.05 ± 0.01 | NA |
| C<sub>eff</sub> binding peptide (μM)<sup>e</sup> | 2.35 | ND | 500 | NA |
| Relative Brightness<sup>f</sup> | 2.37 | ND | 1.16 | 0.26 |
| Time constant (s)<sup>g</sup> | 36 | ND | 38 | NA |
| ND, not determined. NA, not applicable. |  |  |  |  |
| <sup>a</sup> Taken from (36). |  |  |  |  |
| <sup>b</sup> All negative contrast, except for ePDZblock-0.6 and -GGs (positive contrast). |  |  |  |  |
| <sup>c</sup> Apparent dissociation constant of the ratiometric titration curves in Figure 2b and d. |  |  |  |  |
| <sup>d</sup> Hill equation parameter obtained by fitting the ratiometric titration curves from Figure 2b and d. |  |  |  |  |
| <sup>e</sup> Calculated using equations 13 and 15 in Supplementary Note S1. For binding peptide 480, we used the ELISA IC<sub>50</sub> value from (36) instead of an actual K<sub>D</sub>. This probably causes an overestimation of C<sub>eff</sub> for ePDZblock-480 and ePDZblock-480-T. |  |  |  |  |
| <sup>f</sup> In absence of 0.6 peptide. Excitation with 488 nm. ePDZblock-480 was chosen as the reference. |  |  |  |  |
| <sup>g</sup> Time constant of the ratiometric fluorescence decay after 0.6 target peptide addition (Supplementary Figure S8). |  |  |  |  |

We next optimized the connective linkers within the sensor. ePDZblock-480 was chosen as the starting template due to its higher apparent sensor affinity and larger excitation-ratiometric response. The ePDZ-cpFP linker was optimized first by screening 149 variants in bacterial lysates (Figure 2c), resulting in an improved variant in which the cpFP-binding peptide linker was then optimized by similarly screening 166 variants. We then selected the top three best-performing variants in terms of Δ*R /R* and fluorescence and expressed these in HeLa cells. The construct with linkers LA and TP was observed to have largest ratiometric dynamic range *in cellulo*. We selected this variant for all further experiments and named it ePDZblock-480-T. Some variants with a positive contrast but comparatively small dynamic range were discovered as well, but were not pursued for further optimization. This also showed that a lysate screen of only 315 colonies can significantly improve NanoBlock biosensor responses.

Titrations of ePDZblock-480-T and its corresponding control constructs were performed as before (Figure 2d and Supplementary Figure S6b), confirming the improved sensor response. Interestingly, the construct incorporating the 0.6 (high-affinity) binding peptide did respond to the addition of free 0.6 peptide, with an even higher dynamic range than ePDZblock-480-T (Table 2), even though its unoptimized equivalent did not respond. We hypothesize that the altered linkers lower the effective concentration *C*_eff_ of binding peptide 0.6 (Supplementary Note S1 and Table 2), enabling its displacement by free 0.6 peptide at high concentrations. As expected, ePDZblock-GGS-T did not respond to the target peptide.

Additional *in vitro* characterization revealed similar absorbance and fluorescence spectra as for the ALFABlocks (Figure 2e-f), as well as a pKa close to 7 (Supplementary Figure S7), similar to other sensors based on cpFPs. Moreover, the ePDZblocks respond to their target peptide within a minute (Supplementary Figure S8 and Table 2).

### ePDZblocks allow for FACS sorting of live cells

We next expressed ePDZblock-480-T in HeLa cells. To verify the biosensor response to the target peptide, we transfected cells with two plasmids simultaneously: one encoding the sensor, and a second one encoding an mCherry variant engineered to be non-fluorescent (Figure 3a). This ‘dark’ mCherry could furthermore be expressed in isolation or fused to the 0.6 target peptide. We also incorporated a HaloTag protein domain after an IRES sequence so that we could clearly identify doubly-transfected cells during data analysis.

**Figure 3:**
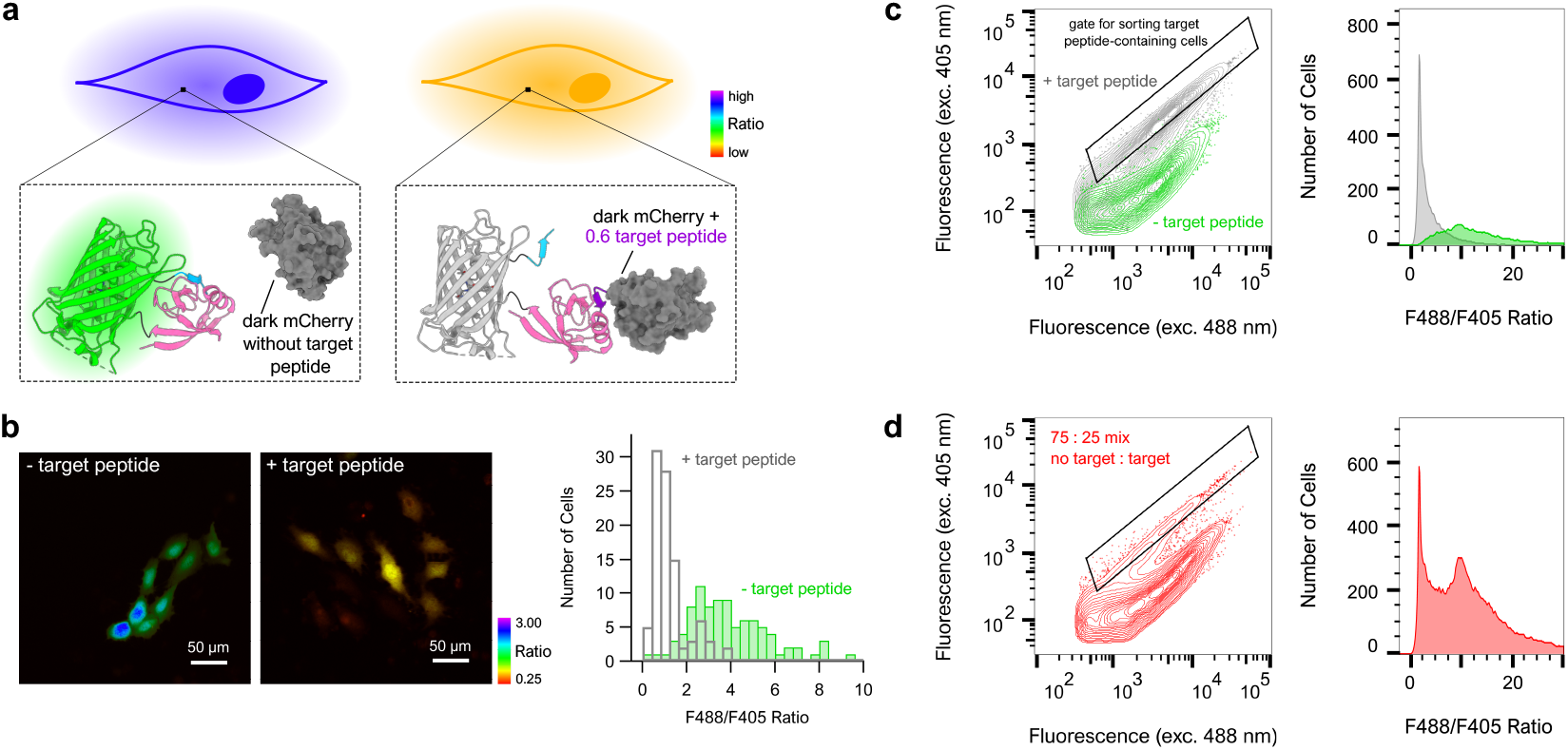
ePDZblocks respond to the 0.6 target peptide in mammalian cells. **a)** Schematic representation of the experimental setup. **b)** Widefield fluorescence microscopy images of HeLa cells doubly transfected with dark mCherry and ePDZblock-480-T. The dark mCherry is either untagged (left image), or tagged with the 0.6 peptide (right image). The color scale depicts the per-pixel F488/F405 ratio. The histogram depicts the ratio distribution for cells without (n=96) and with target peptide (n=95). Outliers in the histogram removed, see Supplementary Figure S9e for the full histogram. **c-d)** Flow cytometric analysis of HeLa cells doubly transfected with dark mCherry and ePDZblock-480-T. The dark mCherry is either untagged or tagged with the 0.6 peptide. In **d**, a mix of untagged (75%) and tagged (25%) dark mCherry is analyzed. Left, a contour plot showing fluorescence intensity after excitation with 488 nm- and 405 nm-light. A gate that could be used for sorting target peptide-containing cells is indicated. Right, histograms of the excitation ratio distributions of the analysed cells. Outliers with a F488/F405 ratio higher than 30 excluded.

Figure 3b shows example images recorded on cells expressing ePDZblock-480-T and the untagged and tagged mCherry. We found that ePDZblock-480-T expressed in the cytosol as expected, and did not form the punctate structures encountered for the ALFAblocks. Comparison of the excitation-ratiometric fluorescence microscopy images acquired on cells expressing peptide-tagged and untagged dark mCherry revealed a Δ*R /R* of 2.86 (negative contrast, Figure 3b and Supplementary Figure S9e). The ePDZblock-GGS-T control sensor did not respond to the target peptide (Supplementary Figure S9b, d) while ePDZblock-0.6-T did show a similar response to the target as ePDZblock-480-T (Δ*R /R* = 2.93, negative contrast; Supplementary Figure S9a, c).

Encouraged by the good sensor response observed with fluorescence imaging, we set out to investigate whether we could separate cells with and without target peptide using fluorescence-activated cell sorting (FACS). Such a scenario could arise, for example, when identifying cells that incorporated foreign DNA that expresses (among others) the target peptide. The observed performance was similar to that seen in the microscopy measurements (Figure 3c and Supplementary Figure S10a-b), with a Δ*R /R* of 3.13 for ePDZblock-480-T and 6.97 for ePDZblock-0.6-T (both negative contrast). Furthermore, plotting the ePDZblock-480-T flow cytometry data in a 405 vs. 488 nm-excitation contour plot showed that the two cell populations (with/without target peptide) differed sufficiently to allow selective extraction of only cells expressing the target peptide (Figure 3c). In particular, using the gate indicated in Figure 3c and d, sorting of a 75:25 (non-target:target) mixture would result in a 1.5:98.5 mixture, significantly enriching the target-containing cells (Figure 3d). In contrast, the two populations are not distinguishable in the individual 405 and 488 nm-excitation channels (Supplementary Figure S10c), showing that the excitation-ratiometric readout is a necessary and robust way to distinguish the different cell populations.

## Conclusions

In this research, we have introduced a way to convert an affinity binder such as a nanobody or similar protein moiety into a genetically-encoded, live-cell compatible fluorescent biosensor using our ‘NanoBlock’ design. The NanoBlock concept operates by converting the affinity binder into a sensing domain via the addition of a binding peptide that competes for the paratope of the binder. In principle, such an affinity binder can originate from a broad variety of sources, which we have explored here via the creation of sensors based distinct frameworks, namely a nanobody and a naturally-occurring protein. We have shown that the resulting sensors display a strong fluorescence contrast and that their response can be robustly read out using excitation-ratiometric imaging. We also showed that the performance of these systems can be enhanced by optimizing the linkers connecting the different sensor domains in a comparatively straightforward manner.

The NanoBlock concept requires the presence of a peptide that binds in such a way that it is displaced by the sensor target. Furthermore, the binding affinity of this peptide should be sufficiently low so that it does not (excessively) interfere with the binding of the native target. If the epitope is a linear peptide then suitable peptides can be created by using mutated variants, as we have done in this study. Alternatively, methodologies like phage or yeast display have proven to be capable of generating peptide binders of varying affinities (*39, 40*). More rational binding peptide design could be performed based on structural information or using novel computionanal methodologies for protein and peptide design. Methods such as PepPrCLIP (*41*), RFdiffusion (*42*), ProteinMPNN (*43*) and BindCraft (*44, 45*) could be employed for *de novo* binding peptide development, or even the design of larger binding domains. This peptide development challenge should be further eased by the requirement for a relatively low affinity. The systems created here displayed higher binding affinities than would be required, supporting sensor operation over a broad range of binding affinities.

We derived an analytical model that rationalizes the inverse relationship between NanoBlock affinity and binding peptide affinity, and showed that this relationship can be affected by intramolecular forces and strains depending on binding peptide and linker sequence. Thus, different choices of binding peptides and of linkers can change the overall sensor affinity and dynamic range, offering additional room for optimization. Since the NanoBlock architecture is fully genetically encoded and fluorescent, multiple binding peptides and linker variants can in principle be screened in a straight-forward manner, simplifying the overall design process.

In this contribution, we focused on cpEGFP-containing biosensors. However, the NanoBlock sensing unit offers the versatility to be combined with several other classes of reporting domains. NanoBlocks containing red fluorescent proteins like cpmApple could provide possibilities for multiplexing and more *in vivo*-friendly imaging. Also non FP-based reporters could be explored in the future, including chemigenetic systems like cpHaloTag (*9, 46*) and cpFAST (*47*), which can offer improvements in spectral properties like brightness and photostability (*48*). Existing calcium sensors based on these systems can be used as starting points. Furthermore, non-fluorescent outputs could be considered as well. The implementation of split enzymes could lead to ‘smart’, conditional enzymes that are only active when a stimulus of choice is present.

Apart from proving the overall NanoBlock concept, the ALFA- and ePDZblocks hold intrinsic value for their ability to detect targets containing small peptide tags. We illustrated this using FACS experiments, showing that we could use the ePDZblock excitation-ratiometric signal to sort cells containing the ePDZ target peptide. This offers opportunities in areas such as genomic engineering, where our peptide-targeting NanoBlocks could be used to confirm the presence of tagged knock-in proteins via transient transfection followed by FACS. Due to the small size of the peptide compared to common selection markers like antibiotic resistance genes or fluorescent proteins, such an approach can be expected to increase the efficiency of methods such as CRISPR while also having a reduced influence on the knock-in gene of interest (*49*).

In conclusion, the NanoBlock architecture offers the potential to considerably broaden the range of targets against which fluorescent biosensors can be designed. In doing so, they would make it possible to reveal many more biochemical processes, deepening our possibilities to monitor and understand complex nanoscale systems such as living cells.

## Materials & Methods

### Plasmid construction

For bacterial expression, all NanoBlock constructs were cloned in a pRSET B vector. Construction of the ALFAblock-WT plasmid was done by HiFi DNA assembly (New England Biolabs) of several fragments: PCR-amplified pRSET B, PCR-amplified NbALFA with as a template pET28b-mEGFP-NbALFA (a gift from Motoyuki Hattori, Addgene plasmid #159986), PCR-amplified cpEGFP and linkers with as a template GCaMP6s (an in-house plasmid), and synthethic DNA encoding the ALFA-tag. All binding peptide variants of this construct were created through a modified QuikChange protocol (Agilent) or through site-directed mutagenesis using the KLD enzyme mix (New England Biolabs).

Construction of pRSET B-ePDZblock-480 was done by HiFi DNA assembly of PCR-amplified pRSET B and synthetic DNA encoding ePDZblock-480. All binding peptide variants of this construct were created through site-directed mutagenesis using a modified HiFi DNA assembly protocol.

For mammalian expression, NanoBlock constructs were subcloned in pcDNA3.1 using HiFi DNA assembly. Dark mCherry (mCherry Y72N), untagged or tagged with a target peptide, followed by HaloTag behind an IRES sequence was cloned into a pcDNA5/FRT/TO backbone. While this back-bone is designed for use with the Flp-In T-REx system (Invitrogen), it acts as a regular pcDNA3.1 backbone when transfected into unmodified mammalian lab cell lines, which is how we used it. The dark mCherry-IRES-HaloTag plasmid was constructed using HiFi DNA assembly of PCR-amplified fragments with in-house plasmids as templates. ALFA-tag (N-terminal) and ePDZ 0.6 peptide (C-terminal) were added through Golden Gate assembly with the ALFA-tag encoded on synthetic DNA and site-directed mutagenesis using the KLD enzyme mix, respectively.

All primers and synthetic DNA were ordered from Integrated DNA Technologies and all constructs were verified by DNA sequencing (LGC Genomics, Eurofins, GENEWIZ or Plasmidsaurus).

### Protein purification and *in vitro* characterization

NanoBlock genes were all expressed from the pRSET B vector, which contains an N-terminal 6xHis tag.

ALFAblocks were expressed in SHuffle T7 *E. coli* cells (New England Biolabs). Cultures were grown in Terrific Broth (TB) at 30 °C until an OD_600 *nm*_ of around 0.5 was reached, whereafter they were transfered to 16 °C for further overnight expression. ePDZblocks were expressed in JM109(DE3) *E. coli* cells (Promega). Cultures were grown similarly as for the ALFAblocks, but the initial temperature was 37 °C and 0.5 mM of IPTG was added when switching to 16 °C.

Pellets were resuspended in TN buffer (100 mM Tris-HCl, 300 mM NaCl, pH 7.4) with 20 mM imidazole and incubated with lysozyme and a cOmplete Mini, EDTA-free protease inhibitor cocktail tablet (Roche) for 30 minutes at 4 °C. After lysis by sonication (Branson Sonifier 450), the His-tagged NanoBlocks were isolated from the clarified lysate using Protino Ni-NTA agarose beads (Macherey-Nagel). This eluate was further purified by size-exclusion chromatography (SEC) on an ÄKTA Go system and with a HiLoad 16/600 Superdex 75 or 200 pg column or a Superdex 200 Increase 10/300 GL column (all Cytiva), using TN buffer. Fractions were analysed with SDS-PAGE and pooled. Concentrations were determined using absorbance at 280 nm measured with a NanoDrop 2000 spectrophotometer (Thermo Scientific) or using a Pierce BCA protein assay kit (Thermo Scientific). Purified NanoBlock could be stored for several weeks at 4 °C in the dark.

For the *in vitro* characterization experiments, target peptides were purchased from NanoTag Biotechnologies (ALFA peptide) and GenScript (ePDZ 0.6 peptide).

Target peptide titrations were performed by mixing purified NanoBlock and target peptide in a flat black µclear 96-well plate (Greiner) to obtain a NanoBlock concentration of 100 nM or 500 nM and peptide concentrations ranging from 0 to 80 µM. After at least 30 minutes incubation at room temperature, fluorescence was measured in a Tecan Spark platereader (excitation at 405 or 488 nm, emission at 515 nm, bandwidth 10 nm). Titration curves were fitted with a Hill equation.

Fluorescence spectra were obtained with an FP-8550 spectrofluorometer and absorbance spectra with a V-750 spectrophotometer (both Jasco). Excitation spectra were recorded at 525 nm emission, emission spectra were recorded at 405 and 488 nm excitation. Purified NanoBlock (1 µM) was incubated without or with target peptide (400 µM ALFA peptide; 200 µM ePDZ 0.6 peptide) for at least half an hour at room temperature before measuring.

Timelapse measurements of NanoBlock response upon target peptide addition were performed in a Tecan Spark platereader with settings as described above. Purified NanoBlock (100 nM) fluorescence was measured in a kinetic loop. After 5 (ePDZblocks) or 10 (ALFAblocks) cycles, 5 µL of TN buffer or target peptide was manually added at a final concentration of 19 µM (ALFA peptide) or 80 µM (target peptide 0.6). After 5 seconds of shaking, the kinetic loop of fluorescence measuresments was continued. The fluorescence decay after peptide addition was fitted with a single or double exponential function to determine the time constant.

pH titrations were performed in a universal buffer consisting of 50 mM citrate, 50 mM Tris, 50 mM glycine, 100 mM NaCl, and either no or 40 µM target peptide. The pH was adjusted to the desired values, ranging from 4.5 to 10.5. Purified NanoBlock was added at a final concentration of 100 nM (ALFAblock-4) or 50 nM (ePDZblock-480-T). Fluorescence was measured in a Tecan Spark plate reader as described above. Titration curves were fitted with a Hill equation to obtain the pKa.

All data analysis was done using Igor Pro 8 (Wavemetrics) and Microsoft Excel 365.

### Biolayer interferometry (BLI)

The binding kinetics analysis of purified NbALFA (NanoTag Biotechnologies) to the peptide variants (WT, 1, 3, 4, 5) was performed using the Octet RED96e system (Sartorius) applying manufacturer’s recommendations. To this end, 1 µg/mL of biotinylated peptides (with double DoA linker, GenicBio) diluted in Octet buffer (PBS, 0.1% BSA, 0.02% Tween20) were immobilized on streptavidin-coated biosensor tips (SA, Sartorius) and unbound peptides were washed away. For the association step, a dilution series of NbALFA ranging from 1.25 nM – 320 nM was applied followed by dissociation in Octet buffer. Each concentration was normalized to a reference applying Octet buffer for association. Data were analyzed using the Octet Data Analysis HT 12.0 software applying the 1:1 ligand-binding model and global fitting.

### Bacterial lysate screening of linker libraries

Libraries of NanoBlock linker mutants were created in a pRSET B plasmid backbone using HiFi DNA assembly (New England Biolabs). ALFAblock-4 and ePDZblock-480 were used as templates. Linkers were varied in both length (1-3 aa) and amino acid composition, either both linkers at the same time (ALFAblocks) or one linker at a time (ePDZblocks). Amino acid randomization was accomplished using either the 22c-trick (*50*) or NDT codons (*51*). Figure 2c and Supplementary Figure S5a include details on the exact linker composition of the libraries. The ALFAblock library was transformed into SHuffle T7 *E. coli* cells (New England Biolabs) while the ePDZblock libraries were transformed into JM109(DE3) *E. coli* cells (Promega). From the resulting colonies, the most fluorescent ones under blue light were picked and grown in autoinduction medium in 96-deep well plates at 30 °C (SHuffle cells) or 37 °C (JM109 cells) overnight. For the ALFAblock library, 1498 colonies were picked, while for the two ePDZblock libraries, 149 and 166 colonies were picked. After storing a sample of each culture as a 10% glycerol stock at -80 °C for sequencing purposes, the cell pellets were either stored at -20 °C or directly lysed using a 50/50 mixture of B-PER (Thermo Scientific) and TN buffer. The NanoBlock sensor activity of 100 µL of cleared lysate was tested in a timelapse experiment in a microplate reader, for up to 20 minutes. The ALFAblock library was screened with a BioTek Cytation 5 plate reader (excitation at 488 nm, emission at 515 nm, bandwidth 10 nm), while for the ePDZblock libraries a Tecan Spark plate reader (excitation at 405 and 488 nm, emission at 515 nm, bandwidth 10 nm) was used. After two or three cycles, the plate reader injector was used to add 5 µL of corresponding target peptide to each well in a final concentration of 19 µM. After 5 seconds of shaking, the kinetic loop of fluorescence measurements was continued.

### Mammalian cell culture and transfection

For microscopy, HeLa cells were cultured in DMEM supplemented with 10% fetal bovine serum (FBS), glutaMAX, and 0.1% gentamicin (all Gibco) at 37 °C and 5% CO2. The HeLa (ATCC-CCL-2) cells were acquired at ATCC and were regularly replaced from frozen stocks. Before transient transfection, cells were seeded in 35-mm glass bottom dishes (MatTek), and allowed to settle overnight. Cells were transfected using the FuGENE 6 Transfection Reagent (Promega) according to the manufacturer’s protocol, at a ratio of 6 µL FuGENE 6 /µg DNA. Cells were transfected with two plasmids simultaneously, the NanoBlock in pcDNA3.1 and the pcDNA5/FRT/TO backbone with target peptide-tagged or untagged dark mCherry and HaloTag separated by an IRES sequence. 1 µg of each plasmid was transfected.

For flow cytometry, HeLa cells were cultured in DMEM supplemented with 10% fetal bovine serum (FBS) and glutamine at 37 °C and 5% CO2. In order to have enough cells for sorting and analysis, sensors were transfected in T25 flasks with 4 µg DNA at a ratio of 4.5 µL FuGENE:DNA. Single color controls were transfected in 6-well plates. After 48 hours, cells were labeled with 200 nM of JF646 for 45 mins and then detached with trypsin. The cells were counted using a Vi-Cell (Beckman Coulter) and the concentrations were standardized across the samples. For mixed populations, cells were then mixed by volume to yield the desired ratios.

### Microscopy

Imaging was performed 48 hours after transient transfection. First, the doubly transfected HeLa cells were stained with 100 nM Janelia Fluor 646 (JF646) HaloTag ligand for an hour at 37 °C and 5% CO_2_. After, cells were washed once and maintained in Hanks’ Balanced Salt Solution with Ca^2+^ and Mg^2+^ (HBSS; Invitrogen). Cells were imaged with an Olympus IX71 inverted microscope equipped with a Spectra X Light Engine (Lumencor), a 10x UplanSApo objective (Olympus), an ORCA-Flash4.0 LT+ sCMOS camera (Hamamatsu), an IX2-RFACA motorized fluorescence cube turret (Olympus), a Lambda 10-B Optical Filter Changer (Sutter) and a H117 high precision motorized stage (Prior). This setup was controlled via custom software based on Igor Pro 9 (Wavemetrics). NanoBlocks were imaged using cyan or violet light for excitation, a ZT488RDC dichroic mirror, and ET525/30 nm and AT515lp emission filters (all Chroma). The JF646-labeled HaloTag was imaged using red excitation light, a HC BS 649 dichroic mirror (Semrock) and an T635lpxr emission filter (Chroma).

### Image analysis

Image analysis and visualization was done using Igor Pro 8 (Wavemetrics) and Fiji. Cells were manually selected in the 488 nm-excitation channel, after which cells with no signal in the JF646 channel were excluded to ensure that all analysed cells contained both plasmids. After background subtraction, histograms of the average fluorescence intensity per cell and of the F488/F405 ratio were plotted. Median values with and without target peptide were used to calculate dynamic ranges.

### Flow cytometry

Cells were passed through 22 µm filters prior to sorting on a Sony MA900 equipped with 405/488/561/646 nm lasers and a 100 µm sorting chip. Gain in detectors FL-1 (525/50 nm), FL-7 (525/50 from 405 nm excitation), and FL-10 (665/30 nm) was set using non-transfected and single color controls. Results were gated and plotted with FlowJo 10.10.0 software. Side and forward scatter were used to exclude debris, dead cells and cell aggregates. The 488 nm-excitation and JF646 channels were used to select doubly transfected cells. Cells with a low signal after 405 nm-excitation were excluded as well. The full gating strategy can be found in Supplementary Figure S11. Median values with and without target peptide were used to calculate dynamic ranges.

## Data Availability

Data used as part of this manuscript will be made available on Zenodo upon acceptance.

## Acknowledgements

We are grateful to NanoTag Biotechnologies and Steffen Frey for providing us with low-affinity ALFA-like peptide sequences and advice concerning the use of NbALFA and the ALFA-tag. We thank Luke Lavis (Janelia Research Campus) for providing the JF646 dye. We thank Juliann Tyler (Janelia Research Campus), Linda Kucun and Oualid Massaoudi for helping with molecular cloning, Damla Temel for helping with the optimization of ALFAblock expression and Iris Govaerts (all KU Leuven) for mammalian cell culture maintenance. We would also like to thank Daria Ezeriņa and Joris Messens (both VIB-VUB) for gifting us the SHuffle T7 *E. coli* cells and Hideaki Mizuno (KU Leuven) for providing access to equipment. We thank Robert E. Campbell and Yusuke Nasu (both University of Tokyo) for advice and insightful discussion concerning the bacterial lysate screens, Brandán Pedre (KU Leuven) for valuable suggestions related to the excitation-ratiometric read-out, and Serge Muyldermans (VUB) for insightful discussion related to the NanoBlock concept and the use of nanobodies. We are also grateful to Hana Valenta, Wim Vandenberg (both KU Leuven) and Karine Breckpot (VUB) for helpful suggestions and discussion. SD and VVD thank the Research Foundation Flanders (FWO Vlaanderen) for their fellowships (1190424N and 12B2Y24N). AGT is funded by the Howard Hughes Medical Institute (HHMI). The work in the lab of AGT was supported by HHMI through the Janelia Visiting Scientist Program. We acknowledge support from the Research Foundation-Flanders through grants G090819N and G010723N, and the KU Leuven via C14/22/088.

## Author Contributions

PD conceived the project with input from AGT. PD and AGT supervised research. SD, VVD, AGT and PD designed experiments. SD, AGT and VVD performed experiments. SD analyzed the data. DIF performed the BLI measurements under supervision of UR. PD developed the analytical model of NanoBlock binding affinity with input from SI. SI contributed critical discussion. SD and PD wrote the manuscript with input from all authors.

## Competing Interests

The authors declare no competing interests.

## Materials & Correspondence

Correspondence and requests for materials should be addressed to Peter Dedecker or Alison G. Tebo. All plasmids encoding the biosensors developed in this work will be made available via Addgene upon acceptance.

## Notes

### Competing Interest Statement

The authors have declared no competing interest.

