## Supplementary information for "NanoBlocks: creating fluorescent biosensors from affinity binders using competitive binding"

October 27, 2025

#### Supplementary Note 1: Binding affinity of a NanoBlock sensor

We created an analytical model of the NanoBlock sensors in order to better understand how the presence of the binding peptide affects the overall sensor affinity. We denote the target of the NanoBlock (the sensed stimulus) as $T$ and the complex of NanoBlock and target as $C$. When it is not bound to the target, the affinity binder inside the NanoBlock can be bound to the binding peptide ($B$) or it can be free of any bound partner ($F$):

$$\mathrm{B} \rightleftharpoons \mathrm{F} \underset{-\mathrm{T}}{\overset{+\mathrm{T}}{\rightleftharpoons}} \mathrm{C} \tag{1}$$

We define $K_{\mathrm{bp}}$ as the dissociation constant for the equilibrium $B$ and $F$ (binding of the binding peptide to the affinity binder), while $K_{\mathrm{bd}}$ describes the dissociation of $T$ from the NanoBlock-target complex $C$, resulting in free $T$ and $F$ (binding of the target to the affinity binder). In other words, $K_{\mathrm{bp}}$ describes how easily the binding peptide can associate with the affinity binder, and $K_{\mathrm{bd}}$ describes how easily the affinity binder can associate with its natural target. The equilibrium expressions for these reactions are given by

$$K_{\mathrm{bp}} = \frac{F}{B} \tag{2}$$

$$K_{\mathrm{bd}} = \frac{FT}{C} \tag{3}$$

where the symbols denote the equilibrium concentrations of the species involved. We also have mass balances relating the total sensor concentration $C_0$ and target concentration $T_0$:

$$C_0 = C + B + F \tag{4}$$

$$T_0 = T + C \tag{5}$$

Using equations (2) to (5), we wish to obtain an expression for the apparent dissociation constant $K'_D$ of a NanoBlock sensor for its target:

$$\mathrm{B} \underset{-\mathrm{T}}{\overset{+\mathrm{T}}{\rightleftharpoons}} \mathrm{C} \tag{6}$$

This system of equations can be solved analytically to obtain the fractional concentrations of each state at equilibrium:

$$\frac{B}{C_0} = \frac{K_{\text{bp}}(C_0 - T_0) - K_{\text{bd}}(K_{\text{bp}} + 1) + D}{2K_{\text{bp}}(K_{\text{bp}} + 1)} \tag{7}$$

$$\frac{C}{C_0} = \frac{C_0 K_{\text{bp}} + K_{\text{bd}} K_{\text{bp}} + K_{\text{bd}} + K_{\text{bp}} T_0 - D}{2K_{\text{bp}}} \tag{8}$$

$$\frac{T}{C_0} = -\frac{C_0 K_{\text{bp}} + K_{\text{bd}} K_{\text{bp}} + K_{\text{bd}} - K_{\text{bp}} T_0 - D}{2K_{\text{bp}}} \tag{9}$$

$$\frac{F}{C_0} = \frac{K_{\text{bp}}(C_0 - T_0) - K_{\text{bd}}(K_{\text{bp}} + 1) + D}{2(K_{\text{bp}} + 1)} \tag{10}$$

where

$$D = \sqrt{(-C_0 K_{\text{bp}} + K_{\text{bd}} K_{\text{bp}} + K_{\text{bd}} + K_{\text{bp}} T_0)^2 + 4C_0 K_{\text{bd}} K_{\text{bp}}(K_{\text{bp}} + 1)} \tag{11}$$

In titrations, one is typically concerned with determining the total target concentration $T_{0;50\%}$ required in order for half of the sensor to be bound to the target. We can obtain this by setting the left-hand side of equation (8) to 1/2 and solving for $T_0$:

$$T_{0;50\%} = \frac{C_0}{2} + \frac{K_{\text{bd}}}{K_{\text{bp}}} + K_{\text{bd}} \tag{12}$$

The apparent dissociation constant $K'_D$ of the system is equal to $T_{0;50\%}$ when $C_0$ is sufficiently small, so that

$$K'_D = K_{\text{bd}}\left(1 + \frac{1}{K_{\text{bp}}}\right) \tag{13}$$

If there is no binding peptide then $K_{\text{bp}} = \infty$ and $K'_D = K_{\text{bd}}$, as expected. Overall, we find that the apparent dissociation constant for the NanoBlock is the dissociation constant of the used affinity binder multiplied by a correction factor equal to $1 + 1/K_{\text{bp}}$.

What is the meaning of $K_{\text{bp}}$? Our result can be viewed within the pharmacological framework that considers binding of a molecule to a target in the presence of a competitive agonist A that is freely present in solution (*1–3*). Since the agonist competes for the binding site, its effect is to reduce the apparent affinity of the binding, resulting in an effective dissociation constant $K'_D$ equal to

$$K'_D = K_{\text{bd}}\left(1 + \frac{A}{K_A}\right) \tag{14}$$

If we consider the binding peptide to be such a competitive agonist then we can combine equations (13) and (14) into

$$\frac{1}{K_{\text{bp}}} = \frac{C_{\text{eff}}}{K_{\text{pep}}} \tag{15}$$

where $C_{\text{eff}}$ is the effective concentration of the binding peptide and $K_{\text{pep}}$ is the dissociation constant of the binding peptide and affinity binder when they are both free in solution (that is, not fused in an intramolecular construct).

Within this framework, $C_{\text{eff}}$ collects all of the changes in binding that arise due to the integration of the components into a unimolecular construct, including steric and other intermolecular constraints in the fusion construct. $C_{\text{eff}}$ is equal to the concentration of sensor ($C_0$) when binding of the binding peptide and the affinity binder is equally likely to occur within the same sensor molecule as between different sensor molecules. Higher values for $C_{\text{eff}}$ indicate that the binding (predominantly) happens via intramolecular association. A higher $C_{\text{eff}}$ therefore reflects reduced steric hindrances to intramolecular association within the same sensor molecule.

### Supplementary Figures

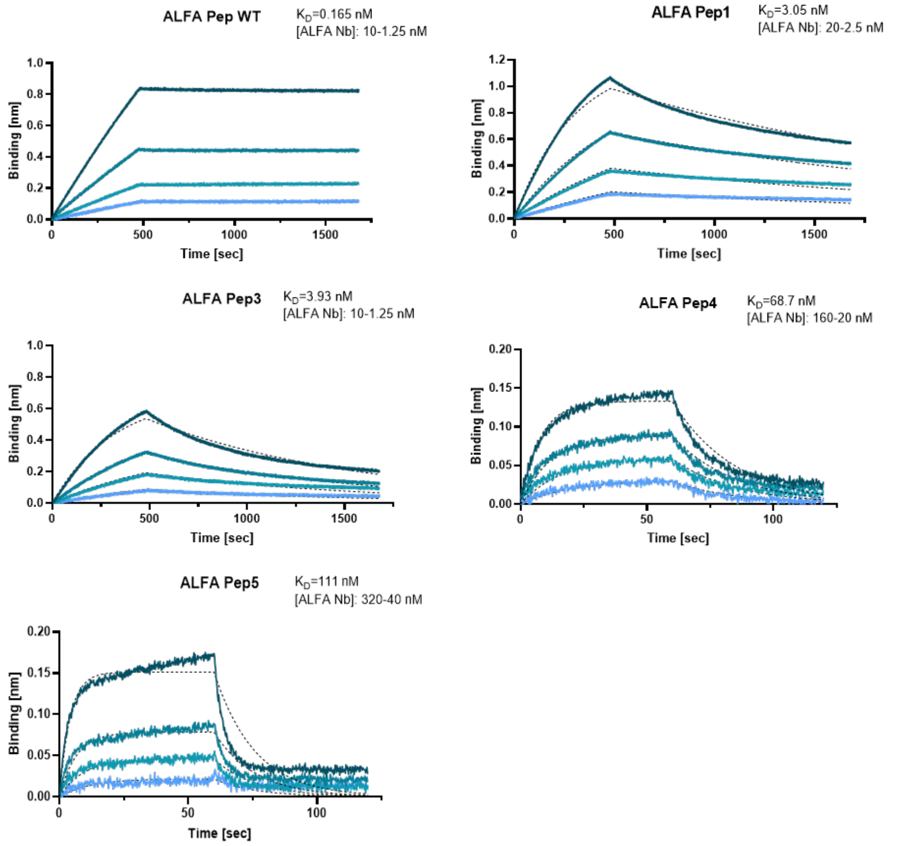


Figure S1: **Binding kinetics analysis of the interaction between NbALFA and the ALFA binding peptides with biolayer interferometry.** The shown sensorgrams depict the association and dissociation of NbALFA to immobilized biotinylated ALFA-like peptides. NbALFA was applied at four different concentrations (shown in different colors), ranging between the indicated values.

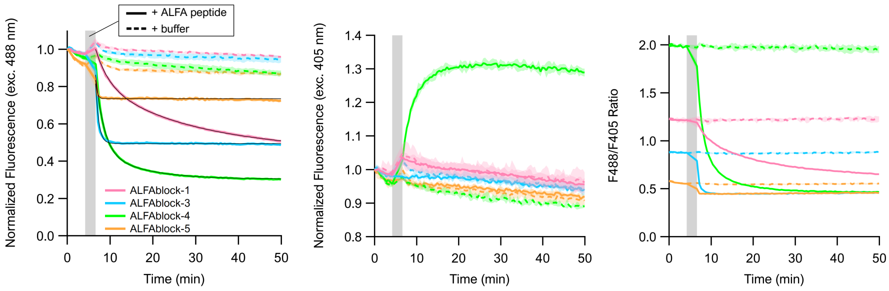


Figure S2: **Response times of the ALFAblocks to the addition of free ALFA peptide.** Time-lapse measurements of ALFAblocks containing different binding peptides, before and after addition of 19 µM ALFA-tag peptide (solid lines) or buffer (dashed lines). The shaded gray area indicates the time needed for peptide/buffer addition to all measured wells. Colored dashed and solid lines represent mean ± s.d., n=3. Black lines are single (ALFAblock-3 and -5) or double (-1 and -4) exponential fits. Left, 488 nm-excitation. Middle, 405 nm-excitation. Right, F488/F405 excitation ratio. Purified sensor at 100 nM.

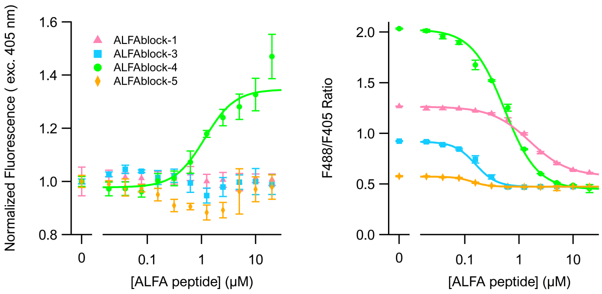


Figure S3: **Excitation-ratiometric behavior of the ALFAblocks.** ALFA-tag peptide titration curves for ALFAblocks with different binding peptides. Solid lines represent Hill equation fits, data points represent mean ± s.d., n=3. Left, 405 nm-excitation. Right, F488/F405 excitation ratio. Corresponding 488 nm-excitation curves can be found in Figure 1b. Purified sensor at 100 nM.

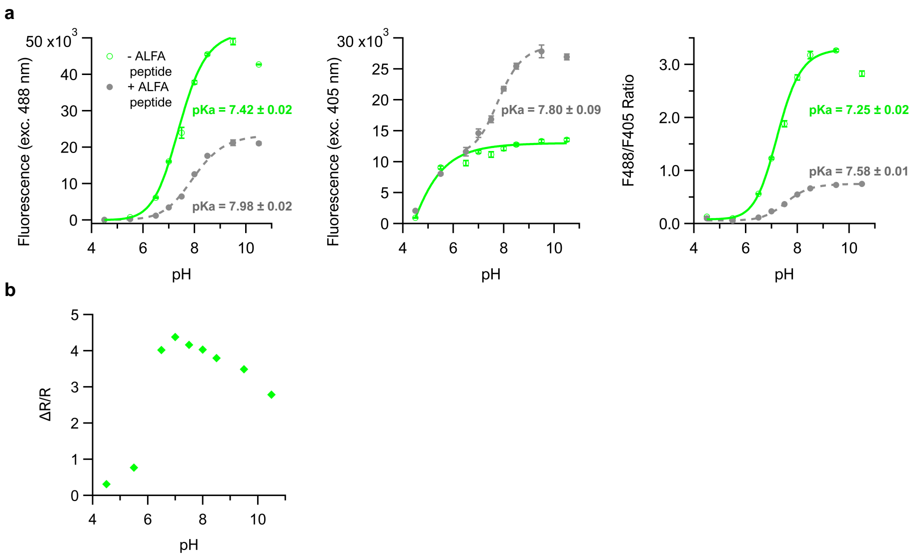


Figure S4: **pH sensitivity of ALFAblock-4. a)** pH titration curves of ALFAblock-4. Dashed (+ ALFA peptide) and solid (- ALFA peptide) lines represent Hill equation fits, data points represent mean ± s.d., n=3. Left, 488 nm-excitation. Middle, 405 nm-excitation. Right, F488/F405 excitation ratio. Purified sensor at 100 nM. **b)** Ratiometric dynamic range ($\Delta R/R$, negative contrast) versus pH.

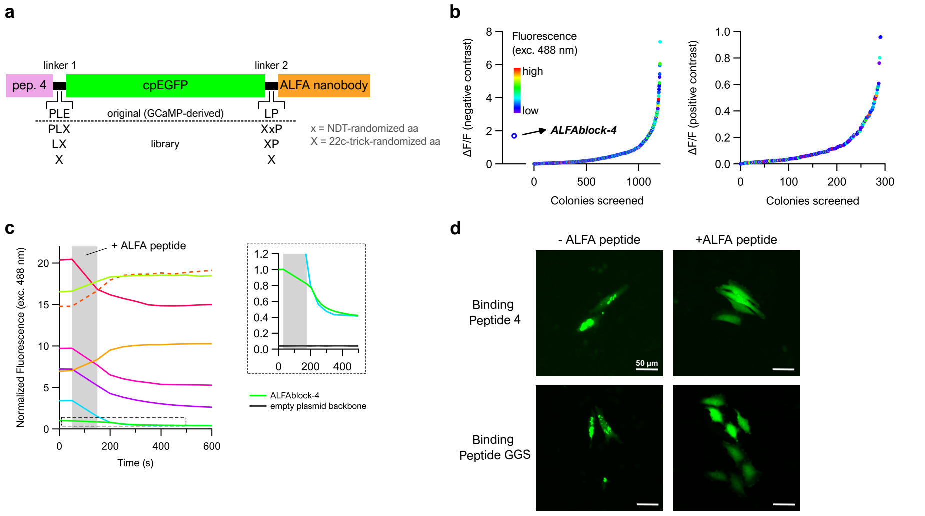


Figure S5: **ALFAblock-4 linker optimization and mammalian cell expression. a)** Linker optimization of ALFAblock-4 through bacterial lysate screens. Both linkers were optimized simultaneously, through creation of a library varying in linker length and amino acid composition. **b)** Results of the ALFAblock-4 linker lysate screen. Each dot represents one screened linker variant, ranked by increasing dynamic range (exc. 488 nm). Left, variants with a negative contrast upon ALFA peptide addition. Right, positive contrast. The color code indicates the fluorescence intensity when excited with 488 nm-light. A total of 1498 colonies was screened. **c)** Timelapse curves of selected ALFAblock-4 linker variants before and after addition of 19 µM ALFA peptide. The shaded gray area indicates the time needed for ALFA peptide addition to all measured wells. Each curve corresponds to the bacterial lysate of one linker variant. **d)** Widefield fluorescence microcopy images of HeLa cells transfected with a linker-optimized ALFAblock-4 (top, dashed curve in **c**) or its corresponding GGS-binding peptide control (bottom). These ALFAblocks were cotransfected with eiter an untagged dark mCherry (left) or an ALFA-tagged dark mCherry (right). Excitation with 488 nm.

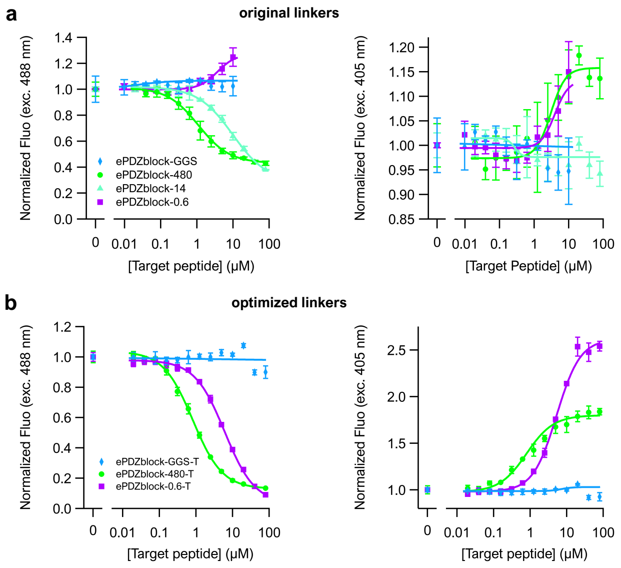


Figure S6: **Excitation-ratiometric behavior of the ePDZblocks.** Target peptide (0.6) titration curves of ePDZblocks with different binding peptides. Shown is the normalized fluorescence upon excitation with 488 nm-light (left) and 405 nm-light (right). The corresponding F488/F405 curves can be found in Figure 2b and d. Sensors either contain the original, GCaMP-derived linkers **(a)** or the optimized linkers from the bacterial lysate screen **(b)**. Solid lines represent Hill equation fits, data points mean ± s.d., n=3. Purified sensor at 100 nM.

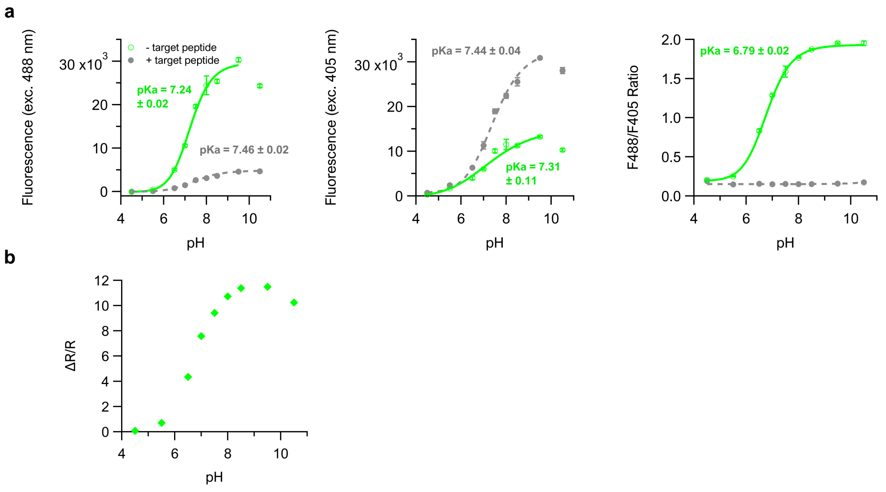


Figure S7: **pH sensitivity of ePDZblock-480-T. a)** pH titration curves of ePDZblock-480-T. Dashed (+ target peptide 0.6) and solid (- target peptide) lines represent Hill equation fits, data points represent mean ± s.d., n=3. Left, 488 nm-excitation. Middle, 405 nm-excitation. Right, F488/F405 excitation ratio. Purified sensor at 50 nM. **b)** Ratiometric dynamic range ($\Delta R/R$, negative contrast) versus pH.

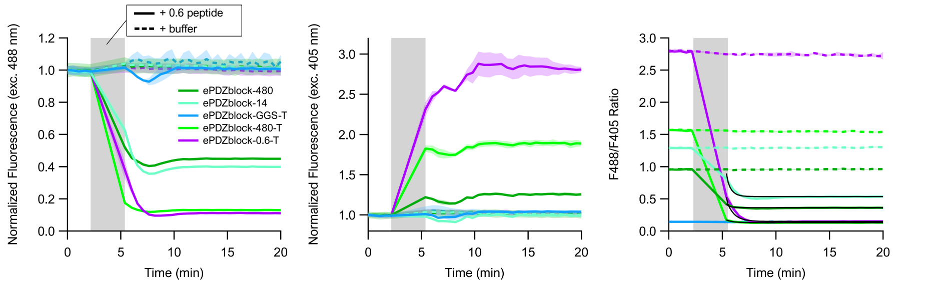


Figure S8: **Response times of the ePDZblocks to the addition of 0.6 target peptide.** Timelapse measurements of ePDZblocks containing different binding peptides and linkers, before and after addition of 80 μM target peptide 0.6 (solid lines) or buffer (dashed lines). The shaded gray area indicates the time needed for peptide/buffer addition to all measured wells. Colored dashed and solid lines represent mean ± s.d., n=3. Black lines are single (ePDZblock-480 and -480-T) or double (-14 and -0.6-T) exponential fits. Left, 488 nm-excitation. Middle, 405 nm-excitation. Right, F488/F405 excitation ratio. Purified sensor at 100 nM.

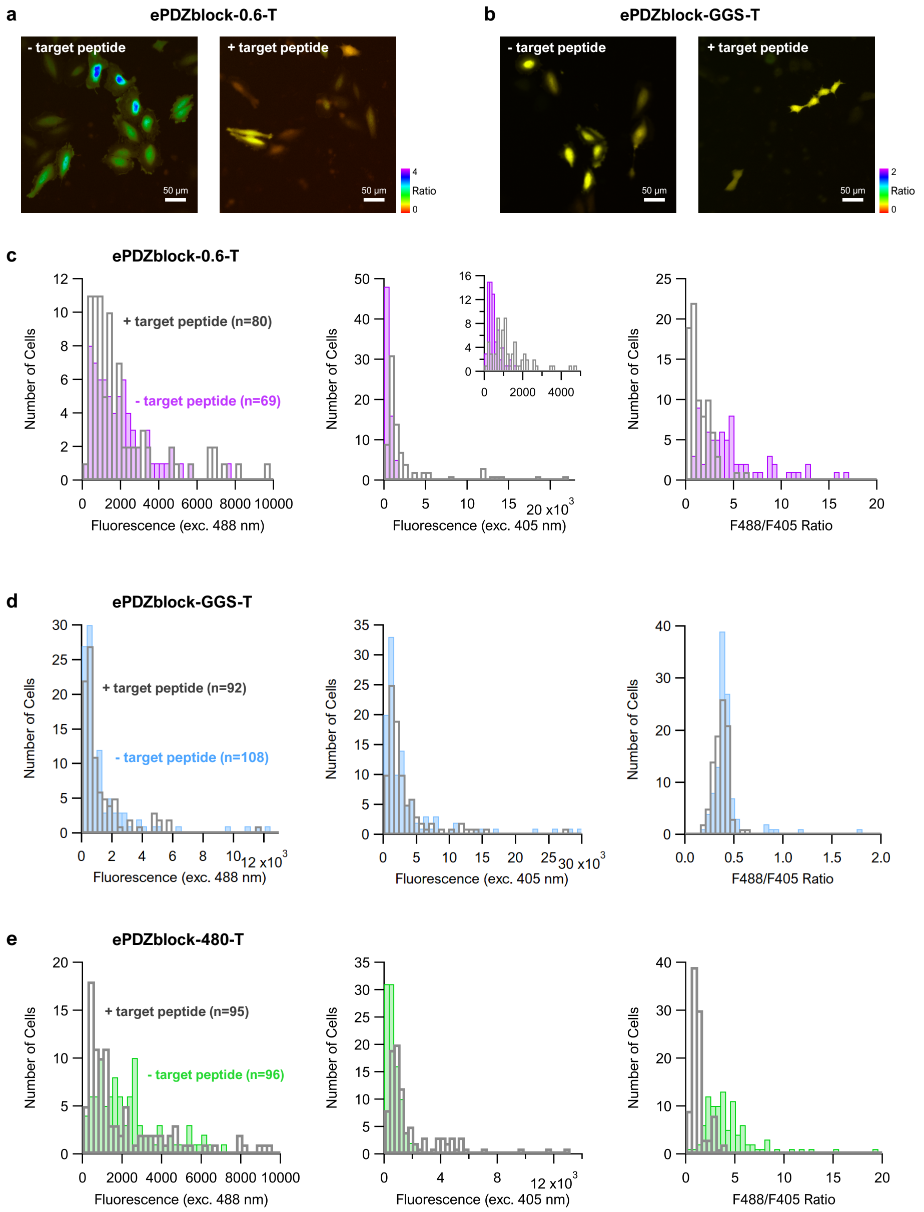


Figure S9: **Widefield fluorescence microscopy of HeLa cells transfected with ePDZblock-T sensors. a, b)** Fluorescence microscopy images of HeLa cells doubly transfected with dark mCherry and ePDZblock-0.6-T **(a)** or ePDZblock-GGS-T **(b)**. The dark mCherry is either untagged (left image), or tagged with the 0.6 peptide (right image). The color scale depicts the per-pixel F488/F405 ratio. **c, d, e)** Histograms depicting the distribution of fluorescence intensity upon 488 (left) and 405 nm-excitation (middle) and of the F488/F405 ratio (right) for cells without and with target peptide. Inset in **c** is a zoomed-in portion of the histogram. Subpanel **e** corresponds to the data in Figure 3b.

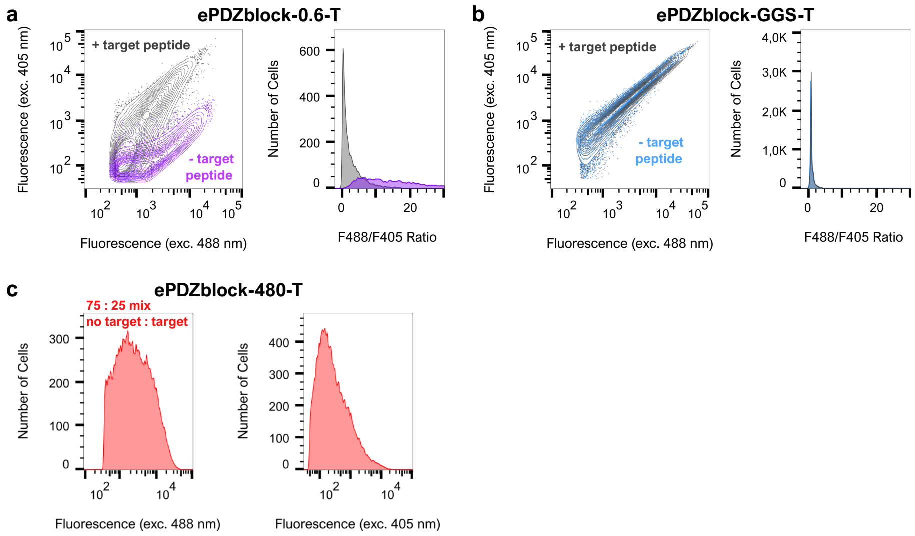


Figure S10: **Flow Cytometry of HeLa cells transfected with ePDZblock-T sensors. a, b)** Cells were doubly transfected with ePDZblock-0.6-T **(a)** or ePDZblock-GGS-T **(b)** and dark mCherry. The dark mCherry is either untagged (- target peptide), or tagged with the 0.6 peptide (+ target peptide). Left, a contour plot showing fluorescence intensity after excitation with 488 nm- and 405 nm-light. Right, histograms of the excitation ratio distributions of the analysed cells. Outliers with a F488/F405 ratio higher than 30 excluded. **c)** Fluorescence intensity histograms of cells doubly transfected with ePDZblock-480-T and dark mCherry. A mix of untagged (75%) and tagged (25%) dark mCherry was analyzed. Left, excitation with 488 nm. Right, excitation with 405 nm. Outliers with a F488/F405 ratio higher than 30 excluded.

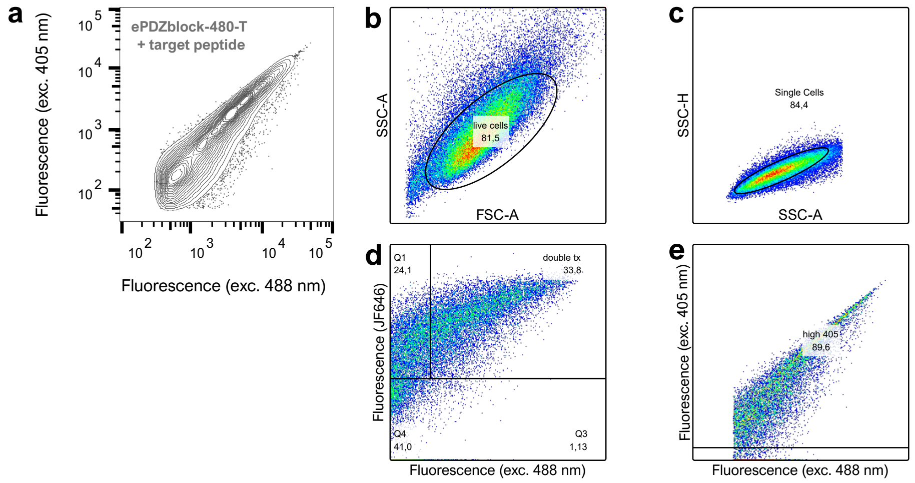


Figure S11: **Flow cytometry gating strategy.** The used gates were the same for all samples, and are illustrated here for HeLa cells transfected with ePDZblock-480-T and target peptide-tagged dark mCherry (**a**). The plasmid encoding dark mCherry also contains HaloTag after an IRES sequence, which was stained with JF646. First, cell debris was excluded (**b**, side vs. forward scatter). Then, single cells were selected using gate **c** (side scatter height vs. area) and doubly transfected cells were selected with the top right gate in **d** (JF646 fluorescence vs. sensor fluorescence upon 488 nm excitation). Finally, cells with too little sensor fluorescence upon 405 nm excitation were excluded using gate **e** (fluorescence upon 405 nm vs. 488 nm excitation).
